# Disparate estimates of intrinsic productivity for Antarctic krill across small spatial scales, under a rapidly changing ocean

**DOI:** 10.1101/2025.03.27.645857

**Authors:** M Mardones, E.T. Jarvis Mason, F. Santa Cruz, G. Watters, C.A. Cárdenas

## Abstract

Understanding the spatio-temporal dynamics of Antarctic krill (*Euphausia superba*) productivity along the Western Antarctic Peninsula (WAP) requires the application of robust analytical approaches. Both design-based and model-based methodologies have been employed to address this challenge. The Spawning Potential Ratio (SPR) provides valuable insights about population indicators. In this study, we analyzed the spatial and temporal variability of the SPR for Antarctic krill by applying a Length-Based Spawning Potential Ratio (LBSPR) model to 20 years of fishery-dependent length composition data. Results showed spatial and temporal heterogeneity among fishing strata in the WAP, where Gerlache Strait stratum was consistently lower than the 20% SPR reference point, compared with Elephant, Bransfield Strait, South West and Joinville strata. Moreover, we demonstrate the sensitivity of LBSPR to changes in growth parameters, such as *k* and *L_inf_*, which are influenced by environmental variables like chlorophyll. Our findings underscore the necessity of incorporating environmental variability into stock assessment models, such as those based on SPR, to accurately assess krill stock conditions. Given the apparent spatial heterogeneity in intrinsic productivity identified through our SPR estimates, we propose using this approach to establish a management procedure based on a control rule for each stratum. This method adjusts the allocation of catch limits in line with the new management strategy of the Commission for the Conservation of Antarctic Marine Living Resources (CCAMLR). By integrating knowledge about spatial krill dynamics and its intrinsic productivity, advice can be recommended to ensure the sustainable management of krill populations in Subarea 48.1.

## INTRODUCTION

One structural component of the Southern Ocean (SO) is the Antarctic krill (*Euphausia superba*; hereafter krill), a keystone species responsible for efficient energy transfer between lower and higher trophic levels [1,2] within a delicate ecological balance strongly influenced by both environmental and oceanographic conditions [3,4]. Along the SO, the Western Antarctic Peninsula (WAP), is considered one of the most sensitive areas to climate change and has already experienced fast changes in various dimensions [5]. Over the last 40 years, climate driven changes have resulted in warming waters [2,5], declines in seasonal sea ice extent and duration [6,7] and changes in phytoplankton productivity [2,8,9]. WAP is a critical region for krill productivity, serving as a major spawning and recruitment area, and also, as a vital region for breeding and foraging for a number of krill predators [10–13]. Given these characteristics, it is expected that environmental changes in this area have influenced the spatial and temporal dynamics of the krill population [14]. Furthermore, these environmental changes occur at different scales, therefore also implying localised spatio-temporal changes of the krill population within the WAP [15–17].

Krill is the largest marine biomass on Earth [4,10,15] and has been the subject of increasing commercial exploitation over the last 50 years [10,18–21]. Since the early 1990s, the fishery has been concentrated between the WAP and the Scotia Sea (FAO statistical subareas 48.1, 48.2 and 48.3) where 70% of the krill population resides [4,9,10,17,22]. The Commission for the Conservation of Marine Living Resources (CCAMLR) is a decision-making body aimed at conserving Antarctic marine fishing stocks, including krill, and associated ecosystems using the best available science. A specific objective of CCAMLR is minimizing risks associated with harvest rates that may affect the populations and avoiding potential irreversible impacts on the ecosystem [23,24,25,26,27 Art. II].

The krill fishery management scheme establishes catch limits in the Convention Area based on a harvest control rule that depends on two biomass reference points [22,23,25,28]. The first rule (*“the depletion rule”*) is based on the lowest level of harvesting allowed, around 20% of biomass, which could be considered a limit reference point. The second rule (*“the escapement rule”*) is the target level of the fishery, which indicates the statistical distribution of the biomass at the end of the 20-year projection under a constant catch that allows the median escapement of 75% at pre-harvested levels of biomass. This scheme resulted in the set of a regional precautionary catch limit within the WAP area, based on a synoptic estimate of biomass which was established in 1991 [26,29].

Traditionally, fishery management strategies worldwide often use feedback loops where catch limits are adjusted in response to updated stock assessments and significant changes in stock status [13,25,28,30,31]. To achieve this, fishery management entities, including governmental agencies and Regional Fisheries Management Organizations (RFMOs), have established management frameworks centered on Biological Reference Points (BRP). These frameworks are constructed considering the population dynamics of the target species, ecosystem interactions, and the reliability of available data [25,30,32], in order to promote sustainability, not only for the fishery and target stock, but also the surrounding marine communities [30,33]. In this context, CCAMLR has monitoring programs that acquire new and revised krill data; however, it has refrained from using a traditional feedback loop to adjust the operational measures (i.e. precautionary catch limits). This lack of adjustment in management procedure, and the recent expiration, due to lack of consensus of CM 51-07 [34], which distributed catches in space across four subareas (48.1 to 48.4), may compromise the efficacy of krill management and, consequently, krill sustainability. In fact, in Subarea 48.1, where a significant krill fishing activity takes place, concerns remain that even at this spatial scale, the catch limit may be insufficient to prevent localized impacts and indirect effects on other ecological components [3,26]. To address these issues, CCAMLR is undertaking a revision of its management strategy framework and conservation measures for the krill fishery. The challenge is to enhance resource management and conservation by adopting a more precautionary and ecosystem-based fisheries management [4,13,35]. In this new management strategy, the spatial and temporal changes in the krill population and community structure is a relevant issue, where management at local or reduced spatial scales, instead of regional, has been proposed to identify changes and give management advice that considers stock status and overlap with other ecosystem components [13,26,36]. A significant obstacle to achieving small-scale management is the absence of a comprehensive quantitative performance metrics based on BRP to assess the degree to which factors influencing spatial heterogeneity have affected krill populations within the WAP and other regions in the SO [25,26]. Consequently, a paradigm shift is proposed for Antarctic fisheries management, transitioning from the current reductionist framework to a more complex strategy grounded in BRP. This new approach should incorporate spatial considerations and account for both krill population dynamics and ecosystem integrity [22,25,26].

In light of this, understanding the changes in krill population structure becomes crucial, as these shifts, reflected in movement patterns, ontogeny, and biological traits, directly influence the spatial and temporal productivity and intrinsic productivity of krill populations in the WAP [12,37,38]. The intrinsic productivity refers to the ability of a population to reproduce and sustain itself, which depends on reproductive and somatic conditions [39,40]. A common way to estimate it in commercially harvested marine populations, like krill, is to quantify the average number of offspring produced by the population over its lifetime, also known as Spawning Potential Ratio (SPR), and it depends on the growth rate and fishery selectivity [41–47]. SPR can give signals about status in more or less exploited areas, because recruitment is related to overfishing [48,49]. That is, when the reproductive fraction has been severely reduced due to excessive fishing, the success of recruitment and stock replenishment is jeopardised [48,50,51]. The SPR has been widely used due to the advantages related with its easy biological interpretation and also because it uses the length composition of the catches, one of the most abundant and reliable sources of information obtained from fishing activity [41,43,52].

Numerous length-based assessment methods have been used to quantify alterations in intrinsic productivity in harvested populations, specifically the SPR [41,44,47,53]. For example, the Length-Based Spawning Potential Ratio (LBSPR) [44] is a modeling approach to derive SPR and relative fishing pressure estimates from length composition and life history data [43,44,46,54,55,55]. Given that the SPR can be used as an indicator of population status, we can use it to improve our understanding of the degree of spatial heterogeneity of the krill population in the WAP. This may consequently, provide elements for establishing limit and target BRPs and allow for flexible adjustment of catch limits in krill fishery management [49,56]. In addition to fishing, environmental factors play a crucial role in shaping the SPR by affecting essential life history traits of krill [38,57]. Fluctuations in temperature, salinity, and nutrient availability directly affect metabolic rates, growth, and reproductive output [3,58–63,63–65]. In marine environments, elevated temperatures can accelerate metabolism, potentially increasing growth but also demanding greater energy expenditure [8,18,65]. Conversely, nutrient limitation can constrain primary production, reducing food availability and consequently, impacting population growth [8,63].

This study aims to quantify the intrinsic productivity of krill using LBSPR modeling over the past two decades across five strata currently identified by CCAMLR in Subarea 48.1 [28,66]. Additionally, we assess changes in somatic growth, which are influenced by environmental conditions, and test their impact on the intrinsic productivity of the krill population. The results of this spatially explicit, strata-based assessment of productivity using SPR as a key indicator, offer valuable insights to refine and optimize catch limit allocations in WAP. These adjustments are based on krill BRPs to support the sustainable management of the krill fishery in Subarea 48.1, aligning with CCAMLR’s revised fishery management strategy.

## METHODOLOGY

### Study area

The study area corresponds to Subarea 48.1 in the WAP, where most krill fishing activity occurs and where CCAMLR has been working to implement a new management strategy at a reduced spatial scale. To achieve a more precise spatial definition of krill population dynamics and analysis, we considered the five strata proposed by CCAMLR: Bransfield Strait, Elephant Island, Gerlache Strait, Joinville Island, and the Southwest (Fig 1). It is worth noting that while these strata were originally delineated for management planning, they did not initially incorporate biological or population-specific characteristics. Nevertheless, they provide a suitable framework for our analysis.

**Fig 1:**
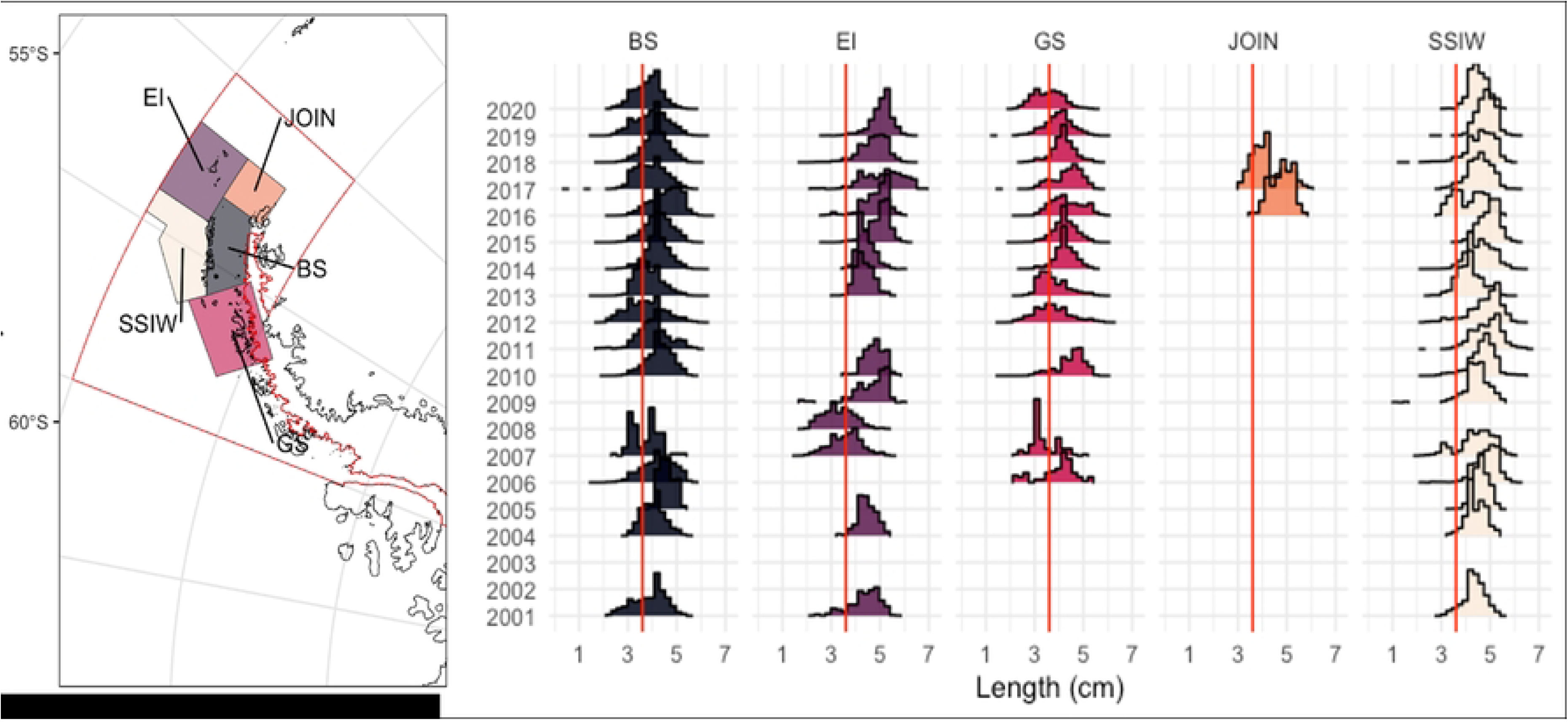
Subarea 48.1 and the WAP management strata considered in the spatio-temporal analysis of intrinsic productivity of krill (BS=Bransfield Strait, EI= Elephant Island, GS= Gerlache Strait, JOIN= Joinville Island, SSWI= South West) and length composition by strata. Red line is length recruit (36 mm)

### Monitoring Data (CCAMLR SISO)

We obtained data from the monitoring of the krill fishery, which has been systematically conducted onboard fishing vessels by scientific observers as part of the CCAMLR Scheme of International Scientific Observation (SISO). Krill length compositions comprising 685,745 individual records were obtained from Subarea 48.1 for the 2000-2020 period, which was grouped by monthly, year and management stratum and grouped into 2 mm bins (Fig 1). The database included information on vessel identification, nationality, georeferenced fishing locations, among other fields. To enhance the representation of size-based population structure for subsequent modeling, descriptive statistics were calculated using the *“tidyverse”* package in R [67,68].

### Environmental drivers of krill growth

To establish environmental drivers of krill growth in the WAP, we focused on the relationships between krill length and three key environmental variables, identified in previous studies as the most influential factors in krill population dynamics [2,8,18,38,69]: 1) Sea Superficial Temperature (SST) using ERA5 monthly mean data from 1991 to present; 2) Sea Ice Index Concentration (SIC) from Nimbus-7 in GeoTIFF format files, from November 1978 to present and 3) Chlorophyll (Chl) data from Bio-Geo-Chemical, L4 (monthly and interpolated) Satellite Observations. We downloaded all data from satellite observation using E.U. Copernicus Marine Service Information (doi.org/10.48670/moi-00021; doi.org/10.48670/moi-00148.) and using native resolution, which represents the finest level of detail captured by the measurement instrument or sampling method. Post-processing techniques, including data cleaning and handling algorithms, manipulation of spatial data, rasterization and projection transformations were facilitated by the *sf* and *ncdf4* packages in R [70,71].

We tested for correlations between krill length compositions from fishery data and environmental variables using a Pearson correlation test. This method is a statistical measure that quantifies the linear relationship between two continuous variables, providing a numerical value, known as the Pearson correlation coefficient (*r*), which indicates both the strength and direction of the linear association between the variables [70].Based on the results of collinearity among variables, the linear models described below were constructed. Also, to analyze the factors influencing krill growth, we used a Generalized Linear Mixed Model (GLMM) framework [72]. The response variable in the model is *Length*, which serves as a proxy for krill growth. This approach allows us to account for both fixed and random effects, capturing spatial and environmental variability while addressing potential non-independence in the data. The linear model is formulated as follows:

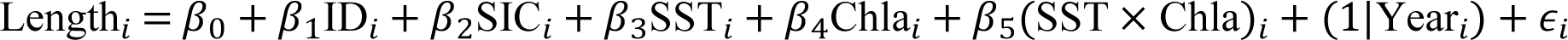

where Length is the krill *Length* for observation *i*, used as a proxy for growth, and β_0_ is the intercept. The fixed effects include ID, which represents the management unit (stratum) as a categorical variable to account for spatial heterogeneity in krill growth. Sea ice concentration (SIC), sea surface temperature (SST), and chlorophyll-a concentration (Chl) are included as environmental covariates, as they influence krill food availability and metabolic rates. An interaction term between SST and Chl is also considered to capture potential synergistic effects on krill growth. A random effect is included for *Year* to account for interannual variability in krill growth. This prevents observed trends from being biased by year-specific factors and mitigates the impacts of data scarcity, which is common across the strata (ID). The residual error, ϵ_*i*_, is assumed to follow a normal distribution. This modeling approach provides a robust framework for assessing krill growth patterns while handling missing data, temporal variability, and potential correlations within the dataset. The estimated slope coefficients and associated 95% confidence intervals (CIs) indicate the strength of the relationship between variables, in which coefficients that overlap zero indicate no relationship with krill length. The variance components indicate the amount of variation that is explained by the random factors. We computed CIs and *p-values* using a Wald *t*-distribution approximation. Along with the mean length, we also configured a model using the 75th percentile of lengths as an alternative way to model the response. In total, six linear models were established, and the selection was based on model performance. Pearson tests and the GLMM were carried out with the *lm4r* and *easystats* packages [72,73]. Analysis and manipulation of variables were grouped into strata using the *sf* and CCAMLRGIS packages [70,74]. A complete description of the linear models used can be found in Supplementary Material 2.

### LBSPR Model

The Length-Based Spawning Potential Ratio (LBSPR) method is a fishery assessment tool that uses the length distribution in a fishery to estimate reproductive characteristics based on life history parameters [44]. It aids in fishery management by offering insights into overfishing risks with minimal data, applicable to both long-lived species with low reproductive output, and short-lived species with highly reproductive outputs associated with a high growth rate, such as krill [75]. LBSPR uses length composition data and assumptions about biological and fisheries parameters to make a rapid assessment of stock status relative to virgin (or unfished) levels.

Previous work has shown that under equilibrium conditions (constant F and no recruitment variability) and assuming a von Bertalanffy growth function, constant natural mortality for all ages, and logistic or jack-knife selectivity, the standardization of the composition of lengths of two populations with the same biological and selectivity parameters are identical [41,44]. The LBSPR model depends on both biological and fishery parameters taken from previous works about krill life history and fishery [75,76], which are described in S1 Table 1. Biological parameters included von Bertalanffy asymptotic length (L_inf_), M/K ratio (natural mortality and von Bertalanffy K in a coefficient ratio), length at 50% maturity (L_50_), length at 95% maturity (L_95_), Coefficient of Variation of length (CVL), and maturity by length (maturity ogive). Fishery parameters include length at 50% selectivity (SL_50_), length at 95% selectivity (SL_95_), and the F/M ratio (fishing mortality and natural mortality). By fitting the model to krill length data, we can evaluate its performance. Since most estimates of biological and reproduction parameters for krill have been made at the global SO scale [57], we assumed the same set of parameters for the five strata of Subarea 48.1. The methodological development from which the SPR calculation is derived was extracted from [49], where the ratio of lifetime average egg production per recruit (EPR) was calculated for fished and non-fished population. Like any assessment method, the LBSPR model relies on a number of simplifying assumptions. In particular, the LBSPR model assumes population equilibrium and that the length composition data is representative of the harvested population at steady state [43,44], which we assume plausible for krill.

**Table 1:** Antarctic krill SPR estimates from CCAMLR(1) Subarea 48.1, by management strata and year (parentheses represent the standard deviation). BS=Bransfield Strait, EI= Elephant Island, GS= Gerlache Strait, JOIN= Joinville Island, SSWI= South West

| Year | BS | EI | GS | JOIN | SSWI |
| --- | --- | --- | --- | --- | --- |
| 2001 | 0.132 (0.024) | 0.223 (0.062) | - | - | 0.196 (0.011) |
| 2004 | 0.15 (0.008) | 0.219 (0.028) | - | - | 0.21 (0.045) |
| 2005 | 0.25 (0.014) | - | - | - | 0.253 (0.005) |
| 2006 | 0.214 (0.009) | - | 0.152 (0.09) | - | 0.349 (0.025) |
| 2007 | 0.108 (0.078) | 0.095 (0.008) | 0.069 (0.014) | - | 0.277 (0.057) |
| 2008 | - | 0.069 (0.008) | - | - | NA |
| 2009 | - | 0.322 (0.058) | - | - | 0.232 (0.016) |
| 2010 | 0.197 (0.004) | 0.329 (0.02) | 0.242 (0.014) | - | 0.31 (0.008) |
| 2011 | 0.185 (0.012) | - | - | - | 0.426 (0.009) |
| 2012 | 0.156 (0.004) | - | 0.148 (0.003) | - | 0.386 (0.006) |
| 2013 | 0.13 (0.001) | 0.191 (0.013) | 0.123 (0.002) | - | 0.176 (0.005) |
| 2014 | 0.175 (0.002) | 0.201 (0.007) | 0.172 (0.006) | - | 0.311 (0.004) |
| 2015 | 0.19 (0.003) | 0.389 (0.022) | 0.177 (0.003) | - | 0.386 (0.026) |
| 2016 | 0.288 (0.01) | 0.36 (0.057) | 0.293 (0.007) | 0.331 (0.19) | 0.266 (0.033) |
| 2017 | 0.165 (0.004) | - | 0.214 (0.016) | 0.249 (0.022) | 0.27 (0.01) |
| 2018 | 0.165 (0.003) | 0.334 (0.019) | 0.162 (0.004) | - | 0.294 (0.018) |
| 2019 | 0.161 (0.004) | 0.425 (0.02) | 0.141 (0.005) | - | 0.349 (0.006) |
| 2020 | 0.11 (0.004) | - | 0.087 (0.004) | - | 0.24 (0.012) |

<sup>1</sup>Commission for the Conservation of Antarctic Marine Living Resources

### Biological Reference Points for krill

By definition, the SPR is equal to 100% in an unexploited stock, and zero in a non-spawning stock (e.g. all mature fish have been removed, or all females have been caught). The F_40%_, that is, the fishing mortality that allows the escapement of 40% of the biomass to Maximum Sustainable Yield (MSY), is the fishing mortality rate that translates into SPR_40%_, and is used as a BRP for many species worldwide [48,77]. [55] offers insightful considerations regarding BRP and the life history for many species. The study delineates three distinct strategies, denoted as I, II, and III, corresponding to the r and K life history strategies. These strategies are contingent upon varying levels of reproductive productivity (high or low) and the nature of population growth (fast or slow) [78–80]. Krill is an organism with high productivity and relatively rapid growth [75], which corresponds to a Type I species (r-strategy), with M/k ∼ 1 (m=0.4, k =0.43) therefore, it requires a higher SPR for population replenishment [44,55]. These considerations about the life strategy must be taken into account in the design of any fishery management procedure based in SPR. We used two reference points for intrinsic krill productivity, SPR_20%_ as the limit reference point and SPR_75%_ as the target for this fishery, which follows with the 20% and 75% biomass reference points used by the CCAMLR management scheme [22,23,25,28,81].

### Sensitivity scenarios analysis

Environmental factors can exert a significant influence on fish growth rates. Consequently, individual growth trajectories may serve as proxies for the environmental conditions encountered by an organism during its lifespan [59]. SPR estimates from the LBSPR model are susceptible to changes in growth parameters [44,82–86]. Considering this, we tested 10 values between the lower and upper range for *L_inf_* (55 to 65 mm). Secondly, we evaluated three theoretical *k* scenarios related to environmental conditions, expressed as low growth (*k* = 0.2), medium growth (*k*= 0.7, current parameter used in krill management), and high growth (*k*= 1.2) (S1 Fig 2). Each scenario was applied to the five strata within the Subarea 48.1, which comprised a total of 65 combined scenarios. Results were compared with those provided in S1 Table 1 serving as the base model (in particular, *k* = 0.45, *L_inf_* = 60 mm). This analytical approach investigated how fluctuations in environmental drivers, specifically those influencing growth (parameters *L_inf_*and *k*) of organisms like krill, can affect the reproductive success of the species. By examining these interconnections, we aim to elucidate the mechanisms underlying krill population dynamics and potential vulnerabilities in response to future environmental changes. Model equations, figures, tables and auto-reproducible guide to LBSPR model application is available at S1 Appendix.

## RESULTS

### Environmental influences on krill growth

Pearson correlation analysis revealed that all three environmental variables had a statistically significant effect on krill population structure. A strong negative correlation was observed between chlorophyll-a (Chla) and krill length (*r* = -0.43), while an even stronger negative relationship was found between SST and *Chla* (*r* = -0.73) (S2 Fig. 3). Among the evaluated linear models, *Model 5* was identified as the best-performing model, based in metrics such as R^2, RMSE, BIC, and AIC, among others. This model explicitly assessed the effect of spatial strata (ID) on krill length while incorporating key environmental covariates, including sea ice cover, SST, and their interaction with Chl (S2 Fig. 5 and 6, Table 1). Regarding the potential exclusion of the JOIN unit due to its limited sample size (only two years of data), its removal was not necessary, as the linear mixed model effectively accounted for missing data within grouped variables considered as random effects. Some random effects associated with ID were significant, indicating differences among strata. While certain strata exhibited variations in mean length (e.g., GS: β = 4.07, JOIN: β = 0.49), the overall spatial effect remained inconsistent across regions. Additionally, the effect of sea ice was negligible (β = ― 0.0086). To compare the performance of models with collinearity among covariates, we systematically tested models with and without this interaction. Comparisons between models including Chl and SST independently and those incorporating their interaction term suggested a significant influence of this interaction on krill growth variability (β = ― 0.11,*p* < 0.05). Furthermore, to explore alternative response metrics, we evaluated the 75th percentile of length in the *Model L75* model. However, this model demonstrated poor performance and was therefore not included in the final analysis. A ranking of model performance and full description of general linear mixed models used is provided in Table 1 of Supplementary Material 2.

### LBSPR Model Output

LBSPR model outputs exhibited adequate fits, as evidenced by the residual analysis of length composition distributions for krill across the strata (S1 Figs 3 and 4). The model accurately captured the distribution patterns of size classes, indicating its effectiveness in characterizing the population structure and variations in length distribution due to the natural variability in krill populations between years and strata. Comparisons between the simulated length distributions of unfished populations with the observed length distributions within each stratum showed that South West, Gerlache and Elephant Island strata exhibited the greatest differences from the simulated structure. Conversely, the Bransfield Strait and Joinville Island strata demonstrated the empirical data were closer to the virginal condition simulated by the model (S1 Fig 4). Bransfield stratum stood out as having a lower proportion of mature individuals, suggesting a higher prevalence of juveniles. It is important to note that the same maturity parameters were applied across all strata (S1 Fig 5).

Krill SPR varied significantly across years and strata. Notably, none of the strata yielded SPR above the reference level of 75% in any year examined, and all strata, with the exception of Joinville (only 2 years of data) and the last years of the series in Elephant Island, contained years with SPR below the limit reference point of 20% (Fig 2). In fact, intrinsic productivity in Bransfield and Gerlache remained consistently below SPR_20%_ for all years in the time series, while Elephant Island and Joinville Island showed an increasing trend in SPR, which corresponded to the increased prevalence of adults in the catch in recent years (Fig 2). All the estimated values and their associated standard deviation of SPR by stratum and by year are shown in Table 1. The analysis for the whole area 48.1 indicated that SPR values remain stable over time, predominantly below the 0.25 threshold, with no significant trends observed (S1 Table 5).

**Fig 2:**
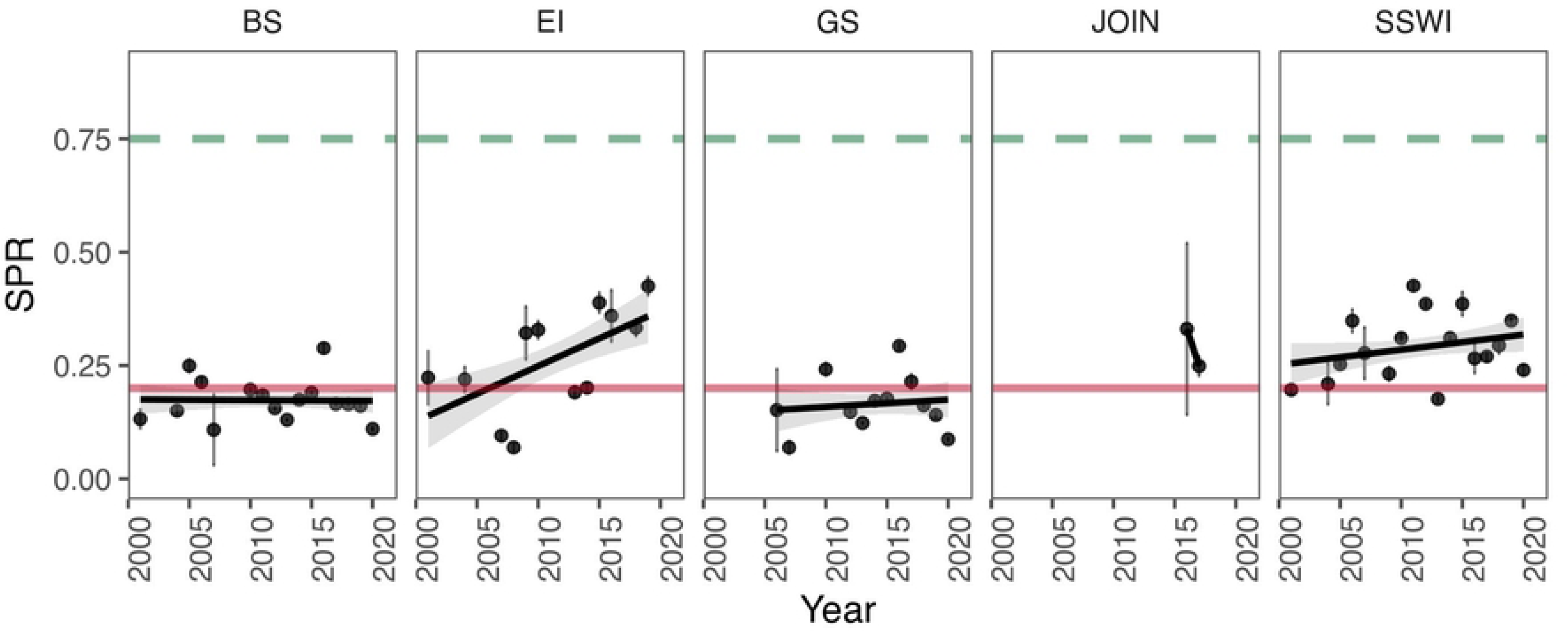
Krill intrinsic productivity (SPR) by management strata and by year. The green dashed line represents 75% SPR (reference target) and the red line is 20% SPR (reference limit). BS=Bransfield Strait, EI= Elephant Island, GS= Gerlache Strait, JOIN= Joinville Island, SSWI= South West.

### Sensitivity analysis

We found that as asymptotic length (L_inf_) increased, the median SPR values generally decreased, with the highest medians observed at L_inf_ 55 mm and the lowest at L_inf_ 65 mm. Specifically, the median SPR values for the Bransfield Strait ranged from 0.28 at L_inf_ 55 mm to 0.12 at L_inf_ 65 mm. In general, lower L_inf_ values in the LBSPR model resulted in higher SPR estimates, a trend that was consistent across all strata. These results suggest a tendency toward greater stability in SPR estimates as L_inf_ increases (Fig 3; Table 2). Regarding the three growth rates (*k*) tested (low, medium, and high), high and medium growth rates produced very low SPR estimates compared to low growth, with particularly high individual growth leading to SPR values very close to the reference level of 75% SPR, while remaining far from the limit reference level of 20% (Fig 4; Table 2).

**Fig 3:**
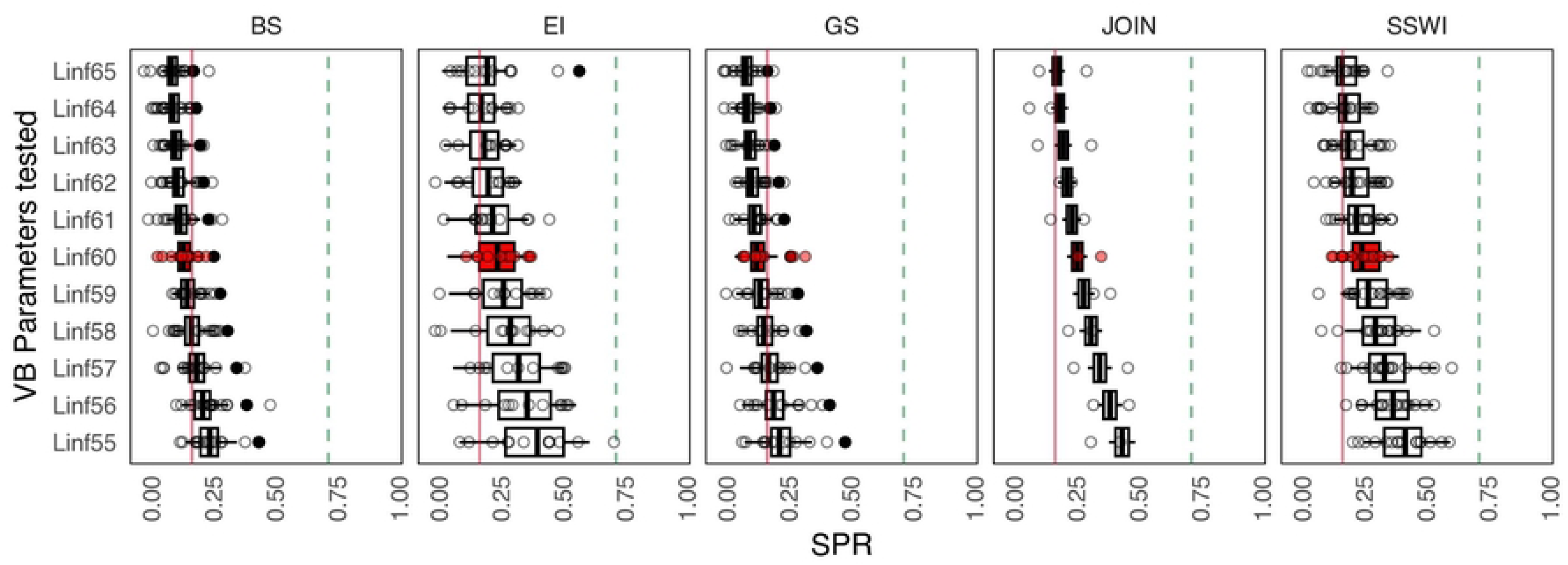
Sensitivity analysis in SPR estimation by management strata in relation to the scenario of a range of VB asymptotic length values for Antarctic krill. The green dashed line represents 75% SPR (reference target), and the red dashed line is 20% SPR (reference limit). Red boxes represent current parameters used in krill management. BS=Bransfield Strait, EI= Elephant Island, GS= Gerlache Strait, JOIN= Joinville Island, SSWI= South West.

**Fig 4:**
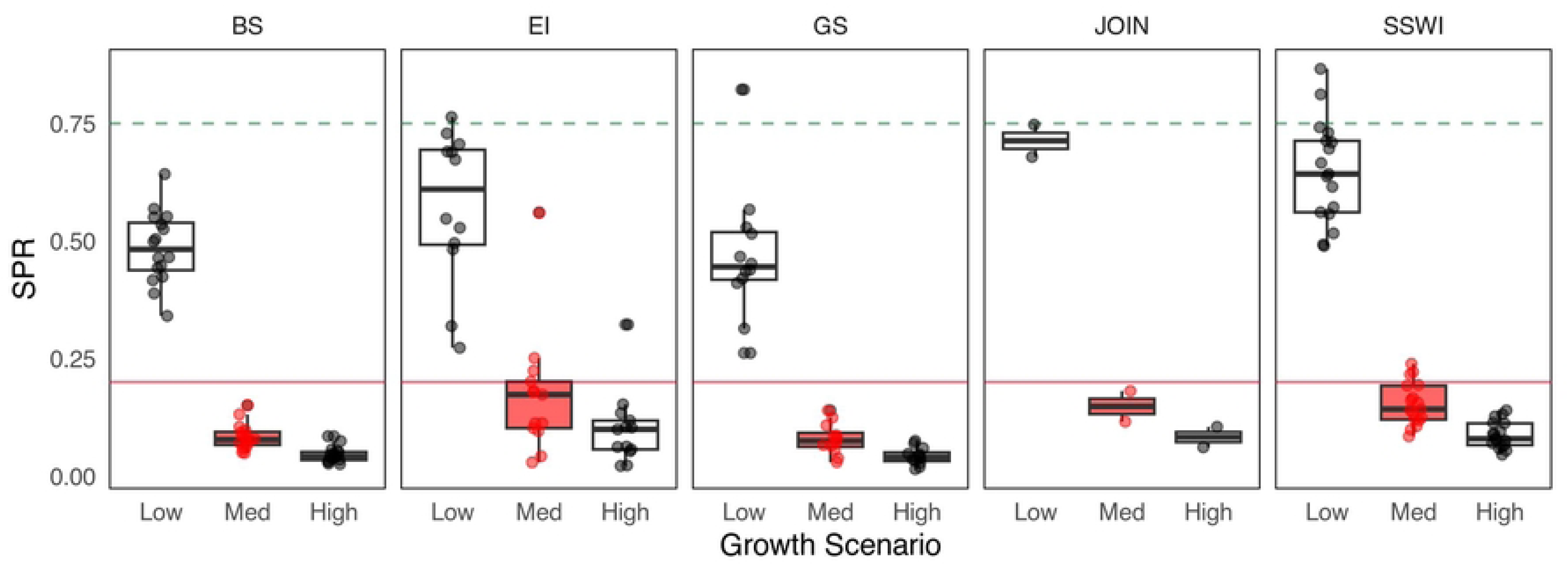
Sensitivity analysis to SPR estimation by management strata in relation to a range of growth coefficients (k) representing three different Antarctic krill growth rates (low = 0.2, medium = 0.7, and high = 1.2). The green dashed line represents 75% SPR (reference target) and the red dashed line is 20% SPR (reference limit). Red boxes represent current parameters used in krill management. BS=Bransfield Strait, EI= Elephant Island, GS= Gerlache Strait, JOIN= Joinville Island, SSWI= South West.

**Table 2:** Antarctic krill LBSPR estimates from CCAMLR Subarea 48.1 by management strata in relation to a range of theoretical krill growth parameters, including eleven asymptotic lengths and three growth rate coefficients (low = 0.2, medium = 0.7, and high = 1.2) (parentheses represent the standard deviation). In bold, the values of the parameters currently used in management (Maschette et al, 2020). BS=Bransfield Strait, EI= Elephant Island, GS= Gerlache Strait, JOIN= Joinville Island, SSWI= South West.

|  |  | BS | EI | GS | JOIN | SSWI |
| --- | --- | --- | --- | --- | --- | --- |
| Asymptotic length ($L_{inf}$) | **Linf55** | 0.28 (0.07) | 0.46 (0.23) | 0.27 (0.11) | 0.47 (0.07) | 0.47 (0.13) |
|  | **Linf56** | 0.25 (0.07) | 0.42 (0.23) | 0.24 (0.09) | 0.42 (0.07) | 0.42 (0.11) |
|  | **Linf57** | 0.23 (0.06) | 0.39 (0.23) | 0.22 (0.08) | 0.38 (0.07) | 0.38 (0.1) |
|  | **Linf58** | 0.21 (0.06) | 0.36 (0.23) | 0.2 (0.08) | 0.35 (0.06) | 0.34 (0.09) |
|  | **Linf59** | 0.19 (0.05) | 0.34 (0.23) | 0.18 (0.07) | 0.32 (0.06) | 0.32 (0.08) |
|  | **Linf60** | **0.17 (0.05)** | **0.32 (0.23)** | **0.17 (0.06)** | **0.29 (0.06)** | **0.29 (0.07)** |
|  | **Linf61** | 0.16 (0.04) | 0.3 (0.21) | 0.15 (0.06) | 0.27 (0.06) | 0.27 (0.07) |
|  | **Linf62** | 0.15 (0.04) | 0.27 (0.18) | 0.14 (0.05) | 0.25 (0.05) | 0.25 (0.06) |
|  | **Linf63** | 0.14 (0.04) | 0.25 (0.17) | 0.13 (0.05) | 0.23 (0.05) | 0.23 (0.06) |
|  | **Linf64** | 0.13 (0.04) | 0.24 (0.15) | 0.13 (0.05) | 0.22 (0.05) | 0.22 (0.05) |
|  | **Linf65** | 0.12 (0.04) | 0.22 (0.14) | 0.12 (0.04) | 0.21 (0.05) | 0.21 (0.05) |
| Growth Scenario (k) | **Low** | 0.49 (0.08) | 0.61 (0.19) | 0.47 (0.14) | 0.71 (0.05) | 0.65 (0.11) |
|  | **Med** | **0.08 (0.03)** | **0.17 (0.13)** | **0.08 (0.03)** | **0.15 (0.05)** | **0.15 (0.05)** |
|  | **High** | 0.05 (0.02) | 0.1 (0.08) | 0.04 (0.02) | 0.08 (0.03) | 0.09 (0.03) |

Sensitivity analysis of the asymptotic length (L_inf_) for Subarea 48.1, shows that SPR estimates consistently fall below the target reference point of 0.75, although some variability occurs across the tested parameter range, with values clustering around the 0.20 threshold. In contrast, sensitivity to the *k* reveals a strong influence on SPR outcomes: the “Low” growth scenario results in higher SPR values (∼0.6), whereas the “Med” and “High” growth scenarios yield significantly lower values, consistently below 0.2. Overall, the analysis points to low spawning potential for krill in area 48.1, with results being robust to L_inf_ but highly sensitive to growth rate assumptions (Fig 5).

**Fig 5:**
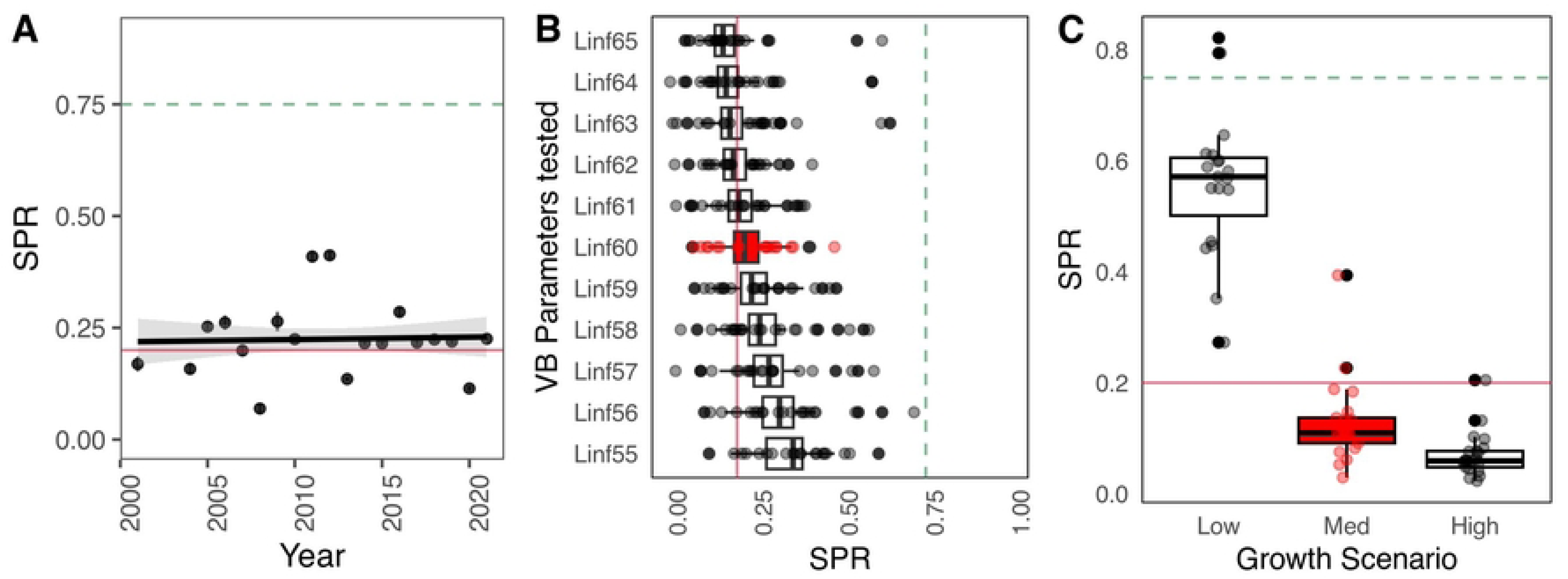
Overall results of SPR estimation and sensitivity scenarios of Antarctic krill for Subarea 48.1. A) SPR values in time series (2000-2020), B) scenario comparisons with VB asymptotic length values and C) SPR variability for growth rates scenario

## DISCUSSION

### Spatial and temporal heterogeneity in intrinsic krill productivity

In most marine populations, spatial heterogeneity in distribution patterns is the rule rather than the exception [87]. Population dynamics and changes in spatial patterns of krill productivity can be influenced by fishing activity, or the fishing activity *“follows”* those changes, concentrating in places where the population is most productive. On the other hand, changes in the ecosystem triggered by environmental and ecological variables are also direct drivers that cause changes in the population structure of krill on the spatial scale [4,17,22]. Consequently, it is imperative to account for spatial heterogeneity in stock structure and distribution when evaluating stock indicators. This is particularly critical for krill, as the fishery has transitioned from a widely dispersed to a highly concentrated spatial distribution into small areas over the past 20 years, particularly in Subarea 48.1. Such concentration may amplify the potential impacts of harvesting. To date, a quantitative assessment of spatial heterogeneity in stock status for krill has been lacking, hindering effective management [4,14,15,26,88].

Our analysis of length composition data from krill fisheries monitoring reveals significant spatial heterogeneity in population structure and its temporal changes. Length composition data from catches were used as they are widely regarded as one of the most representative sources of information on the population dynamics of exploited marine resources [41,46,52]. These data provide valuable insights as an indicator of population status [41], inform biomass estimates [89], and serve as a key component in integrated stock assessment models [90]. Changes in the availability, distribution, and concentration of harvested marine populations such as krill have been reflected in this kind of data, and the use of this type of data and size-based models to estimate the status of the population has been widely used in the world fisheries and allows recommendations for sustainable management [14,41,44,47,50,53].

We verified that the intrinsic productivity of krill varied across management strata and changed over time. Elephant Island and South West Island exhibited the highest SPR levels, showing a positive trend in the latter years of the series. In contrast, the Bransfield Strait and Gerlache strata consistently displayed low SPR levels throughout the entire time series. These strata are characterized by low productivity and a relatively high abundance of immature juvenile individuals, as highlighted in studies by [12] and [38]. Given that krill fishery activity has expanded to these strata, it is imperative to prioritize this area for effective management, especially because it also serves as a recurrent foraging area for krill predators, including baleen whales [91,92]. In Bransfield strata, the SPR has decreased throughout the study period, with the lowest level of 12% SPR occurring in 2020. This may be related to the growing fishing activity that has increased and concentrated in this strata [10,93], which has targeted older krill, thus, reducing reproductive capacity of the population and consequently, decreasing intrinsic productivity. This spatial and temporal heterogeneity of SPR of krill provides a biological mechanism for the consequential changes observed in indicators of productivity yield across the WAP [16,17]. From a holistic point of view, these spatiotemporal differences in krill SPR are likely to significantly impact the maintenance of long-term ecosystem balance and species diversity [4,22]. Higher SPR ensures a larger number of offspring, increasing the chances of population recovery after disturbances or environmental fluctuations, while reduced SPR may cause decreased food availability for dependent predators [20]; hence, fluctuations in intrinsic productivity can have cascading effects on krill population and their predators. Spatial heterogeneity in the population structure of krill (demographic patterns, recruitment) has been widely described for different areas of the SO, specifically in the WAP [11,12,14]. However, these studies are mostly descriptive. In contrast, LBSPR approach is a model-based method that allows us to have a quantitative approximation of the spatio-temporal differences in the krill population structure that can be used to provide meaningful spatially-explicit recommendations for conservation and management, consistent with CCAMLR’s mandate for ecosystem-based fishery management.

Our results support those from previous studies of length-based analysis about growth type [42,46,47], but it is the first time that this type of analysis has been carried out on krill. In this context, two aspects emerge. Firstly, despite recognizing spatial variability in demographic structures among strata, we employed the same set of life history parameters and assumptions for our analysis. Secondly, we identified an information gap regarding biological parameters in krill, crucial for both our analysis and the stock assessment process. Consequently, conducting studies to determine specific life history and maturity parameters by stratum becomes imperative to refine our findings. Nevertheless, sensitivity analysis could serve as a valuable tool to address these existing knowledge gaps.

### Environmental impacts to krill intrinsic productivity

Changes in krill population dynamics are shown in various ways, including distribution, biomass, recruitment and phenology. The main drivers of these changes are related to different environmental variables in the WAP [8,9,14,18,19,21,38,69]. While acknowledging the influence of environmental factors on variables such as biomass and recruitment [12,18,94], it remains imperative to systematically assess spatial and temporal shifts and their impacts on krill intrinsic productivity. Our findings indicate a strong, negative relationship between Chlorophyll concentration and krill size, suggesting that environmental conditions significantly influence krill population structure, which is in accordance with previous research [8,18,21,59,95,96]. Previous research has highlighted the temporal variability in krill population characteristics [12,97]. Similarly, spatial components (strata) and environmental variables, like interaction between Chla and SST contribute meaningfully to explaining length variation in all management strata assessed. Increased chlorophyll levels have been linked to enhanced krill recruitment [6,18], but paradoxically, this can also lead to population declines in mature individuals, consequently reducing SPR according to this study, which is a significant finding. This broader perspective may explain discrepancies with previous analysis. While [96] established a positive correlation between female maturity and krill recruitment, our study expanded this analysis by assessing the spawning potential of the entire population, including both males and females. However, understanding the interplay between fishing activities and environmental conditions on krill physiology and reproduction remains a significant challenge in the face of climate change.

A good practice in the use of stock assessment models, involves testing scenarios to determine the impact of uncertainty sources, like growth parameters, on the output, detects biases and improves reliability [52,98]. Indeed, few studies have performed parameter sensitivity analyses for these methods with real case studies [41,42,44,99]. Given the strong negative influence of chlorophyll on krill growth in the WAP (S2 Fig. 3), we performed krill SPR sensitivity analyses across various values of growth parameters. Recognizing that the parameters of the von Bertalanffy growth curve are correlated [100–102], we applied the analytical derivation by [44] to calculate SPR and independently tested scenarios for each parameter to evaluate their effects.

Testing different growth rate scenarios (*k*) also revealed that slower growth rates (e.g., *k* = 0.2) result in higher SPR values across all strata, indicating greater resilience to environmental and anthropogenic changes (Fig 4). This occurs given that with a smaller *k*, that is, an organism with lower productivity and slower growth [43], there is a higher level of resilience to anthropogenic and environmental changes. Conversely, faster growth rates were associated with lower SPR values, potentially underestimating the risks of overfishing. Overestimating *k* may underestimate SPR, falsely suggesting stock sustainability, while underestimating *k* could lead to overestimated SPR values, providing a false sense of security. These findings underscore the importance of precise, stock-specific estimates of life-history parameters, as misestimations can lead to inappropriate management measures and jeopardize long-term sustainability. [99] and [103] recommend deriving life-history parameters, such as *L_inf_*and *k*, from stock-specific studies to improve the accuracy of length-based assessment methods like LBSPR. In the case of Antarctic krill, while the existing stock assessment model (i.e., the Grym; [76]) incorporates life-history parameters, these data are often outdated. Most research focuses on physiological and reproductive aspects, with limited emphasis on parameters directly used in LBSPR. Addressing these gaps is critical and could involve utilizing available data and estimating missing parameters through bioanalogical methods or Modal Progression Analysis [104]. A quantitative tool like LBSPR, which can detect changes in krill productivity, offers valuable insights into the overall health and resilience of the WAP ecosystem and its krill populations, playing a critical role in informing sustainable management strategies.

### CCAMLR krill fishery management and spatial heterogeneity

A precautionary management approach for the sustainability of commercial krill harvesting relies on maintaining a healthy population through sustainable harvesting practices, which can be influenced by variations in intrinsic productivity. In a standard fishery science context, a robust management procedure typically includes a Harvest Control Rule (HCR), which is based on an algorithm through which the mortality, harvest rate or quota to be implemented for the estimation of the quotas is chosen [30,105]. The HCR provides for adjusting catch limits in accordance with a target fishery reference point (fishing mortality or biomass) that relates to changes in an indicator of stock status, such as SPR. The current krill fishery harvest strategy in Subarea 48.1 is based on a constant catch that is not related to changes in a population and/or ecosystem indicator or associated target fishery reference point, such as Maximum Sustainable Yield (MSY), i.e., maintaining biomass at sustainable levels. CCAMLR has recognized these gaps [13,22,25,26] particularly the absence of a stock assessment process and the lack of consideration for spatial components in the current management recommendations. This procedure was in development over the last 5 years [3,22,28,36] and considered three pivot elements: i) update biomass estimates, whether at macro or small scale; ii) develop a stock assessment approach to estimate precautionary harvest rates or/and quotas; and iii) an overlap analysis for the spatial allocation of catch limits [13,35]. Despite these proposals for assessing krill population dynamics at a finer level, these schemes are not yet operational for decision making [3,26]. In fact, in 2024, the lack of consensus on the implementation of the revised management resulted in the expiration of CM 51-07 and with it, the allocation of catches divided by subareas.

The expiration of CM 51-07 is untimely, as our results only further demonstrate the need for the revised approach and associated implementation of an alternative HCR. The HCR could constitute an *ad hoc* approach for the adjustment of catch limits and, consequently, a scheme for spatial allocation for each stratum within subarea 48.1, based in SPR. With an associated BPR, such as SPR_75%_, this approach provides for a management trigger, such as changes to the harvest rate or adjusting catch limits by strata. The management procedure adjustment presented in this study identified the spatial differences (strata) of SPR, and the proposed HCR could change the harvest level (catch limit) according to the levels of that indicator krill status, which is an adaptation to the rule commonly used and known as a *“hockey stick”* where;

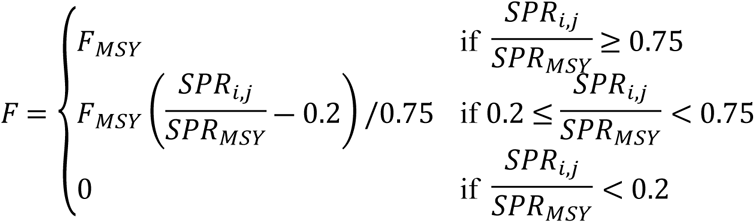

where *i* and *j* indicate the SPR calculated by stratum and year, and SPR_MSY_ represent references in SPR_75%_, which is a common BRP to all stratum. SPR calculation scheme can be easily updated as new data becomes available. The proposal for spatial adjustment and allocation of krill catch limits is not new and has considered different criteria including the catch history, predator demand and krill standing stock by Small Scale Management Unit (SSMU) [24,36,106]. It should be noted that our proposed harvest control rule and spatial allocation scheme is based on an assessment of a single stock of krill, providing a tactical, operational, quantifiable management procedure for krill in Antarctica [30,107]. It also constitutes one of the first biologically-based allocation of spatial catches on krill for the management strata defined in [81].

While our study highlights the utility of the LBSPR method in assessing the intrinsic productivity of krill and its potential as a management tool, it is important to acknowledge the massive data available on krill populations. LBSPR should be viewed as a complementary tool that can provide valuable insights into spatial and temporal variations in krill productivity, especially in the interim period until an integrated stock assessment for krill is fully developed and agreed upon by CCAMLR. This integration of LBSPR into current management practices could bridge gaps in understanding and enhance the effectiveness of krill fishery management, ensuring that decisions are based on the best available science until more comprehensive stock assessment frameworks are established.

## CONCLUSION

This study undertakes the first analysis of the intrinsic productivity of krill in Subarea 48.1 using the quantitative method Length-Based Spawning Potential Ratio (LBSPR) which proved to be a useful tool to assess the influence of multiple stressors on the spatial variability of krill intrinsic productivity while elucidating the environmental drivers of growth and reproductive processes. This approach also provided a robust method to measure temporal trends in spatially-explicit population status relative to limit and target biological reference points. These spatio-temporal variations in intrinsic productivity serve as a fundamental basis as a management procedure, which seeks to integrate the complexities of krill population dynamics into a framework aligned with CCAMLR revised management strategy for Subarea 48.1. The proposed HCR aligns with the revised scheme, and given the absence of biologically-based allocation of catch across strata in the WAP, it would serve well to inform the meaningful allocation of current catch limits in the region, and in turn, contributes to a paradigm shift towards a spatially explicit management scheme. By aligning fishing practices with the natural flows of krill productivity, this approach lays the foundation for a useful coexistence between fishing harvesting and the Antarctic ecosystem, ultimately contributing to the sustainability of this vital element of the Southern Ocean.

## ACKNOWLEDGMENTS

This research was supported by the following funding sources: the INACH “Marine Protected Areas” Program (Grant No. 2409052), the ANID/Millennium Science Initiative Program (Grant No. ICN2021_002), the CCAMLR Scholarship Scheme (2023-2024), and the Doctorate Scholarship from CENTRO-IDEAL at the Universidad Austral de Chile. The authors extend their gratitude to the Secretariat of the Commission for the Conservation of Antarctic Marine Living Resources (CCAMLR) for providing access to the krill fishery data, which was instrumental to this study.

## Author Contributions

- **Conceptualization**: Mauricio Mardones; César Cárdenas, George Watters
- **Formal Analysis**: Mauricio Mardones
- **Methodology**: Mauricio Mardones, César Cárdenas
- **Visualization**: Mauricio Mardones
- **Writing – Original Draft Preparation**: Mauricio Mardones, César Cárdenas, Erica Jarvis Mason
- **Writing– review & editing**: Mauricio Mardones, Erica Jarvis Mason, César Cárdenas, Francisco Santa Cruz, George Watters

## SUPPORTING INFORMATION

- Supporting Information 1.

Key formulas, figures, and code snippets for applying the LBSPR model to study krill populations in the West Antarctic Peninsula (WAP) region can be found at this link: LBSPRKrill.

- Supporting Information 2.

Code and analysis for identifying environmental influences on krill length using correlation and mixed-effects models across spatial and temporal scales can be found at this link: Krill Length Correlation

## Notes

### Competing Interest Statement

The authors have declared no competing interest.

